# Long-term fasting remodels gut microbial metabolism and host metabolism

**DOI:** 10.1101/2024.04.19.590209

**Authors:** QR Ducarmon, F Grundler, C Giannopoulou, A Loumé, N Karcher, M Larralde, S Romano, MR MacArthur, SJ Mitchell, F Wilhelmi de Toledo, G Zeller, R Mesnage

**Affiliations:** Structural and Computational Biology Unit, European Molecular Biology Laboratory, Meyerhofstr. 1, 69117 Heidelberg, Germany; Buchinger Wilhelmi Clinic, Wilhelmi-Beck-Straße 27, 88662 Überlingen, Germany; University Clinics of Dental Medicine (CUMD). Division of Regenerative Dentistry and Periodontology. 1 Michel Servet str, 1211 Geneva, Switzerland; Leiden University Center for Infectious Diseases (LUCID), Leiden University Medical Center (LUMC), Albinusdreef 2, 2333 ZA Leiden, Netherlands; Lewis-Sigler Institute for Integrative Genomics, Princeton University, Princeton NJ, USA; Ludwig Princeton Branch, Ludwig Institute for Cancer Research, Princeton University, Princeton NJ, USA; Department of Nutritional Sciences, School of Life Course Sciences, Faculty of Life Sciences and Medicine, King’s College London, SE1 9NH London, UK

## Abstract

Long-term fasting has become a promising research subject for its potential of treating and preventing metabolic diseases. However, little is known about its impact on the functional capacity of the gut microbiome and the combined effect on the serum metabolome. Here, we demonstrate extensive remodelling of the gut microbial ecosystem in humans (n=92) after an average of 9.8 days of fasting (∼250 kcal / day). Fasting transiently affected the relative abundance of the majority of bacterial species (306 decreased and 210 increased out of 772). Species changes could largely be explained by their genomic repertoire of carbohydrate-active enzymes (CAZymes), which were investigated here for the first time. Fasting induced extensive abundance changes in CAZyme families, depleting families with dietary fibre substrates and increasing families with host-derived glycan substrates. Likewise, we observed extensive changes in the serum metabolome, with 382 out of 721 metabolites significantly affected (246 increased and 136 decreased). In-depth metagenome-metabolome co-variation analysis suggested *Oscillibacter* species to be key producers of indole-3-propionic acid, a crucial metabolite for cardiometabolic health. Together, our results provide an unprecedented view on the impact of long-term fasting on gut microbiome composition and function.

## Main

Caloric restriction and fasting regimes have recently become popular interventions for improving cardiometabolic health, with the potential of preventing or even curing a variety of cardiometabolic diseases^1–4^. The intestinal ecosystem has been suggested as a key source of these beneficial therapeutic effects^5^. However, scientific evidence for a direct implication of the gut microbiome in (cardiometabolic) health effects of fasting is limited to animal studies^6,7^. While causality is difficult to address in fasting humans, we even lack well-powered high-resolution studies of the gut microbiome and its functional potential in this context.

The fasting microbiome represents an underexplored aspect of our health, which is relevant both for therapeutic fasting interventions and for basic physiology, since the gut microbiome exhibits cyclic changes in line with circadian eating^8,9^. The impact of fasting on bacterial (genus) composition is well-characterised, and bacteria believed to be capable of utilising host-derived substrates to proliferate at the expense of bacteria relying on dietary substrates have been reported before^5^. However, remodelling of the gut microbiome at the resolution of species and metabolic pathways including complex carbohydrate metabolism remains largely elusive.

In this study, we provide a high-resolution view on the effects of long-term fasting on the human faecal microbiome through the lens of metagenome functional changes and link these with host and microbial metabolism. By employing carbohydrate-active enzyme (CAZyme) analyses, we uncover pronounced differences between many CAZyme families likely reflecting a broad substrate switch from dietary fibre utilisation to host-derived glycan utilisation. Furthermore, whether a gut bacterial species increases or decreases in abundance during fasting can largely be predicted from its genomically encoded CAZymes. Serum metabolomics confirmed expected enrichments of ketone bodies directly after fasting, but additionally revealed more unexpected changes in microbial metabolites, including a depletion of indole-3-propionic acid (IPA). Integration of serum metabolomics with gut metagenomics data coupled to bacterial genome analysis uncovered *Oscillibacter* species as putatively novel, abundant IPA producers in the human gut.

## Methods

### Study Design

Here we analysed data from two studies: The OralFast observational trial with 40 participants and the GENESIS trial with 58 subjects, all of whom fasted at the Buchinger Wilhelmi Clinic (BWC) in Überlingen. The protocol of the OralFast study was approved by the Ethical Committee of the State Medical Association of Baden-Württemberg (F-2022-025 on 1.06.2022) and registered in clinicaltrials.gov (NCT05449249). Subjects were recruited at the Buchinger Wilhelmi clinic in Überlingen, Germany in September 2021. The GENESIS study is a prospective, monocentric, single-arm interventional study. The study protocol for the GENESIS study was approved by the medical council of Baden-Württemberg on October 7, 2021. It was registered on clinicalTrials.gov (NCT05031598) and recruitment took place between August 2021 and June 2022.

### Participants

Healthy subjects over 18 years of age who registered for a stay of more than 10 days at the BWC and completed the fasting programme could participate in the OralFast cohort. The presence of a contraindication to fasting like cachexia, anorexia nervosa, advanced kidney, liver or cerebrovascular insufficiency, dementia or other severely debilitating cognitive disease and pregnancy or lactation period led to exclusion. We further excluded participants who smoked, took antibiotics within the last 8 weeks, as well as the intake of probiotics within the last 4 weeks, or had periodontal treatment in the last 6 months. The GENESIS targeted women and men aged between 20 and 75 years. Inclusion and exclusion criteria were comparable to OralFast and described previously^10^. Fig. 1a shows the selection procedure for both trials.

**Figure 1:**
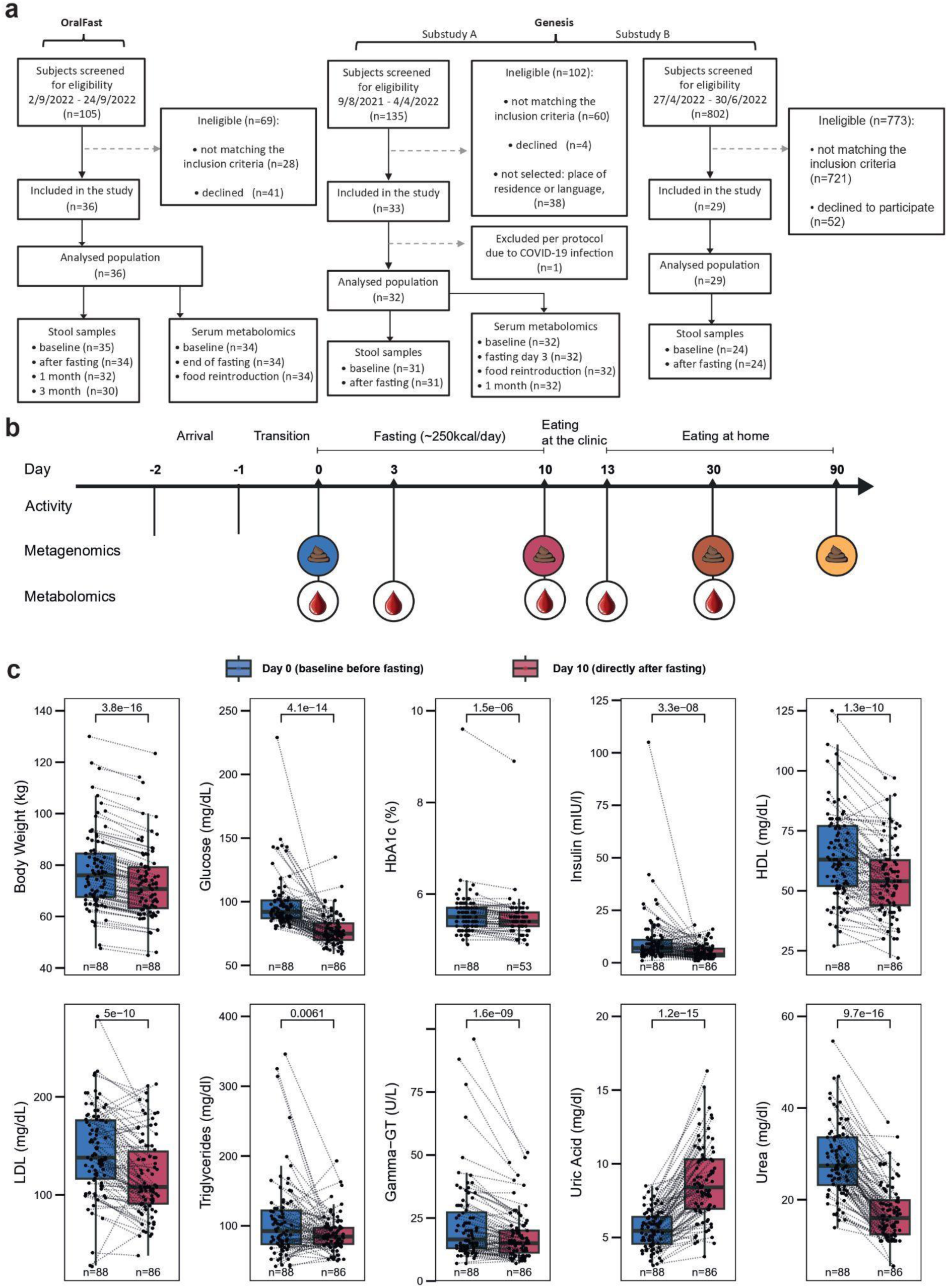
Study design and blood marker changes after fasting. **(a)** Flow chart of study presenting the selection of all included participants. **(b)** Schematic of study design and biosample collection. Participants arrived at the clinic one to two days prior to initiation of the fasting regimen (day 0). Stool samples for metagenomic shotgun sequencing and serum for metabolomics and clinical markers were collected at indicated time points. Timing of the provided stool sample was somewhat variable, with the first stool sample normally provided between day -1 and 0 and the second stool sample provided at first bowel movement during food reintroduction after 9.8 ± 1.8 days of fasting. **(c)** Changes in clinical blood markers related to host metabolic health and lipid, sugar and urea metabolism. Dotted lines connect the measurements within each participant before and directly after fasting. P-values were obtained using a two-sided paired Wilcoxon test. The boxplot centre value corresponds to the median, the box indicates the interquartile range, and whiskers extend to 1.5 times the interquartile range.

### Fasting programme

All subjects underwent a medically supervised long-term fasting program according to peer-reviewed guidelines^11^. In brief, the fasting was minimally supplemented with energy intake limited to 200–250 kcal/day, consisting of 0.25 L of freshly squeezed fruit juice at noon, 0.25 L of vegetable soup in the evening, and 20 g of honey. A gradual reintroduction of a plant-based diet took place over three to four consecutive days, with caloric intake ranging from 800 to 1,600 kcal/day.

### Laboratory examinations

Anthropometric measurements were performed by trained nurses every morning according to the BWC standards. Body weight was assessed while participants were lightly dressed using a Seca 704 stadiometer (Seca, Hamburg, Germany). Blood pressure and heart rate were measured once on the non-dominant arm in sitting position after a 5-minute rest using a boso Carat professional device (BOSCH+SOHN GmbH u. Co. KG, Jungingen, Germany). Blood samples were drawn by trained medical-technical assistants in the morning between 07:30 and 09:30 h. The baseline sample collection was conducted on the transition day or the first day of fasting in the beginning of the stay at BWC. Another blood sample was collected at the end of the fasting period. Additional examinations which were conducted on the GENESIS cohort are described elsewhere^10^. The analysis of the laboratory parameters in Germany at the MVZ Ravensburg was described in detail previously^12^.

### DNA extraction of faecal samples

Faecal samples were self-collected by the patients using the EasySampler Stool Collection Kit (GP Medical Devices, Holstebro, Denmark). The collection tubes contained DNA/RNA Shield™, Zymo Research. Samples were stored at -80°C within 12 hours after collection except for follow-up samples which were sent by post. The samples were processed and analysed with the Microbiome Analysis Service: Shotgun Metagenomic Sequencing (Zymo Research Europe, Freiburg, Germany). DNA was extracted using the ZymoBIOMICS®-96 MagBead DNA Kit (Zymo Research, Irvine, CA) according to the manufacturer’s instructions.

### Metagenomic shotgun sequencing

Metagenomic DNA samples were profiled using shotgun metagenomic sequencing. Sequencing libraries were prepared with the Illumina DNA Library Prep Kit (Illumina, San Diego, CA) with up to 500 ng DNA input following the manufacturer’s protocol using unique dual-index 10 bp barcodes with Nextera® adapters (Illumina, San Diego, CA). All libraries were pooled in equal abundance. The final pool was quantified using qPCR and TapeStation® (Agilent Technologies, Santa Clara, CA). The final library was sequenced by Zymo Research, Irvine, CA on the NovaSeq 6000® (Illumina, San Diego, CA) platform.

### Metagenomics shotgun sequencing data processing

Raw reads were cleaned as described previously^13^. Taxonomic profiling was performed using mOTUs (v3.1) with default parameters on these quality-filtered reads; species abundance profiles referred to in the following were technically mOTU abundances and species names are followed by mOTUs identifiers in brackets. For functional profiling, reads were further screened for host contamination using kraken2 (v2.1.2) against the human hg38 reference genome with ribosomal sequences masked (Silva_v_138). Functional profiling was performed using gffquant v_2.11 (https://github.com/cschu/gff_quantifier) and a reduced version of the GMGC containing 13,788,251 non-redundant genes^13^. The reads remaining after human-genome filtering were mapped to this reduced human gut catalogue using BWA-MEM v_0.7.17 with default parameters. Only alignments with >45bp alignment length and >97% sequence identity were retained for further analysis. Reads aligning to multiple genes contributed fractional counts towards each hit gene. Alignment counts for a gene were normalised by gene length, then scaled according to the strategy employed by NGLess (https://ngless.embl.de/Functions.html#count) and propagated to the functional features with which the gene is annotated. For KEGG KOs, we retained only KOs of prokaryotic origin according to KOFAMKoala prokaryotic HMMs. Gut microbial modules (GMM) were inferred based on KOs and the R package omixerRpm v_0.3.3 using default parameters^14^. Profiling of carbohydrate-active enzymes was done using Cayman (v0.9.6)^13^.

### External metagenomic dataset usage

To validate and contextualise our findings, we profiled several external (longitudinal) metagenomic datasets with various interventions, including dietary interventions, caloric restriction and antibiotic administration^1,15–18^. For surveying relative abundance and prevalence of IPA-producing species, we used two large-scale external cohorts^19,20^. Prevalence was calculated by first rarefying species counts. All external studies were processed with identical bioinformatic workflows as described above in order to minimise bias due to bioinformatic processing.

### Statistical analysis of metagenomic data

Alpha-diversity was calculated using the estimate_richness function in the phyloseq package. Comparison of alpha diversity for baseline versus other timepoints was statistically tested using a linear mixed model (LMM). Beta-diversity was computed using Bray-Curtis dissimilarity and long-term fluctuation of intra-individual microbiome composition was assessed by comparing each sample to its corresponding baseline sample.

All differential abundance analysis was performed using linear mixed models as implemented in SIAMCAT (v2.5.1)^21^. A random effect was included per participant to control for repeated measures. Prevalence threshold for a feature to be included in differential abundance testing was set at 10%. Pseudocounts differed across the different analyses, with 1e-4 for taxonomic analyses, 0.01 RPKM for CAZy analysis and 1e-7 for GMM analysis. P-values were corrected for multiple testing using the FDR method and adjusted p-values < 0.05 were considered significant. Visualisation of the taxonomic tree was done in iTOL v6^22^ showing only those genera with a prevalence > 40% across baseline and the timepoint directly after fasting. Gene-set enrichment analysis for higher-level CAZy substrates was performed using the fgsea (v1.24.0) package^23^.

### CAZyme annotation of bacterial gut genomes

In order to obtain information on the taxonomic origin of CAZymes, we first annotated high-quality genomes from Almeida et al^24^, which amounted to a total of 101,229 metagenome-assembled genomes (MAGs) and 6,456 isolate genomes, as previously described^13^. We then calculated mean CAZyme family copy numbers and prevalences per species on all genomes belonging to the respective species. Genomic copy numbers and prevalences at genus level were calculated analogously.

### Predicting bacterial abundance change

To predict whether a species would increase or decrease in relative abundance based on genomic CAZyme repertoire, we used genomic CAZyme repertoire information as explained in the “CAZyme annotation of bacterial gut genomes” section. CAZyme genomic copy numbers were used as input features to a binary classification approach predicting whether a species increased or decreased in relative abundance during fasting. We employed Random Forest classifiers (200 trees) and assessed their performance using 10-fold cross-validation with 10 resampling rounds as implemented in the SIAMCAT package and we blocked at taxonomic family level^21^. In order to understand how well we can predict relative abundance change based on bacterial genetic similarity alone, we evaluated a nearest neighbour classifier that assigns abundance change direction via the abundance direction change of the closest bacterial species. In order to obtain genetic distances between species, we computed pairwise hamming distances between amino acid-level GTDB^25^ marker gene sequences obtained from motus 3.1 representative genomes. Marker gene sequences were extracted using the *identify* workflow of the GTDBtk toolkit (v2.1.0 and GTDB DB R207). The multiple sequence alignment of the marker genes was generated using the *align* workflow of the GTDBtk toolkit^26^. The same training and testing folds as for the random forest (blocked by taxonomic family) were used, and each mOTU in the test set was assigned to the closest mOTU in the training set.

### Serum metabolomics measurements

Serum (3uL) was added to 60uL of ice cold 100% methanol, briefly vortexed and extracted on dry ice for 20 minutes. The extract was centrifuged at 21,000xg for 20 minutes at 4C and 20uL of supernatant was added to 80uL of ice cold 100% methanol. A second extraction was performed on dry ice for 20 minutes followed by another centrifugation. 60uL of the supernatant was transferred to snap top microvials for LC-MS analysis.

Metabolite measurements were performed using a Thermo Orbitrap Exploris 480 mass spectrometer coupled to hydrophobic interaction chromatography (HILIC). A Waters XBridge BEH Amide column (150mm x 2.1mm) was used. The gradient used solvent A (95%:5% H2O:acetonitrile with 20mM ammonium acetate, pH 9.4) and solvent B (100% acetonitrile) as follows: 90% B (0.0 to 2.0 min), 90% B to 75% B (2.0 to 3.0 min), 75% B (3.0 to 7.0 min), 75% B to 70% B (7.0 to 8.0 min), 70% B (8.0 to 9.0 min), 70% B to 50% B (9.0 to 10.0 min), 50% B (10.0 to 12.0 min), 50% B to 25% B (12.0 to 13.0 min), 25% B (13.0 to 14.0 min), 25% B to 0.5% B (14.0 to 16.0 min), 0.5% B (16.0 to 20.5 min), then stayed at 90% B for 4.5 min. The flow rate was 150uL/min with an injection volume of 7uL and column temperature of 25C. Electrospray ionization (ESI) source parameters were as follows: spray voltage, 2,300 V (positive) or -2,800 V (negative); sheath gas, 35 arb; aux gas, 10 arb; sweep gas, 0.5 arb; ion transfer tube temperature, 300C; vaporizer temperature, 35C. Data acquisition was performed under a full scan polarity switching modes ranging from 70 to 1,000 m/z.

The LC-MS raw data files (.raw) were converted to mzXML format using ProteoWizard software. EL-MAVEN was used to select peaks based on an internally validated knowns list and generate an output table containing m/z, retention time and intensity for peaks. The default EL-MAVEN parameters were used for peak picking except RT shift was restricted to less than 1 minute and ppm tolerance was set to 10 ppm. Baseline samples of the Genesis study were measured twice to confirm robustness of measurements and these profiles were averaged prior to normalisation. Peak intensities were normalised to the median within-sample peak intensity and log transformed prior to analysis. Serum metabolites that are partially or entirely derived from gut microbes were determined using an isotope tracing strategy.

Time-dependent changes in metabolite abundances were determined by employing linear mixed models using a fixed effect for the run and a random effect for the participant. P-values were corrected using FDR. In order to potentially discover novel associations between bacterial species and serum metabolites (e.g. novel producers of specific compounds) we integrated metagenomic data with serum metabolomic data. To this end, we employed LMMs whereby we modelled metabolite abundance by the microbial abundance with a random effect per participant. As matching data could only be gathered from time point day 0 and day 10 for serum metabolomics with the directly after fasting metagenome sample, a total of 127 matching metagenomes and serum metabolomes were used for this analysis. Only species with a prevalence of at least 30% over all these samples were used for analysis.

### Oscillibacter genome mining

In order to investigate whether *Oscillibacter* species contained the phenyllactate dehydratase gene cluster needed to metabolise tryptophan into indole-3-propionic acid, we extracted the sequence of this cluster from *Clostridium sporogenes ATCC 15579* (ABKW02000002.1, region 317042 to 332438), and downloaded genomes from the mOTUs database (v3.0.1) belonging to *Oscillibacter* mOTUs associated with indole-3-propionic acid changes (*ref_mOTU_v3_04664*, *meta_mOTU_v3_12610*, *meta_mOTU_v3_12282*, *ext_mOTU_v3_18233*). These genomes were then screened using cblaster (v1.3.18)^27^ with DIAMOND (v2.0.13)^28^ set to *sensitive* mode. The presence/absence of each query protein was counted for each genome, and then used to compute the proportion of genomes containing the query proteins within each mOTU.

## Main text

Here we used a large longitudinal sample of participants in two clinical studies and investigated changes in their gut metagenome, serum metabolome and general blood biomarkers (n = 92 individuals, n = 241 shotgun metagenomic samples, n = 230 serum metabolomic samples, Fig. 1a,b). Throughout the manuscript, we used the first stool sample collected directly after fasting (as stool production is severely limited during fasting) to infer changes in microbiome “during fasting”. The fasting protocol was generally well tolerated with no recorded adverse effects, except for a 69-year-old participant who experienced acute ischuria, necessitating treatment with a bladder catheter and tamsulosin. This event was not deemed directly related to the fasting intervention by the study physicians, which allowed for continued participation in the study.

Over the course of the fasting period (9.8 ± 1.8 (mean, s.d.)), body weight of participants was on average reduced by 5.01 +/- 1.59 kg (mean, s.d., see Fig. 1c), in line with findings from previous studies, including ours^12,29^. Energy metabolism of the participants switched from utilising food-derived energy substrates to endogenous sources, as evident from changes in glucose, lipid and urea cycle metabolism (Fig. 1c). This was accompanied by improvements of blood markers which are risk factors for e.g. the development of chronic liver diseases (gamma-glutamyl transferase) and cardiometabolic diseases (triglycerides, lipoprotein cholesterol). In general, improvements were strongest for those individuals with the highest baseline levels of these markers, as was previously shown for blood pressure^30^. All clinical data characteristics for all samples can be found in Table S1 and Table S2.

### Fasting extensively perturbs bacterial species composition in the gut microbiome

To broadly assess fasting-induced changes in the gut microbiome, we first surveyed microbiome composition using a Principal Coordinate Analysis (PCoA) of beta diversity based on Bray-Curtis dissimilarity. This revealed substantial remodelling during fasting compared to baseline, to which microbiome composition returned for long-term follow-up samples after one and three months (Fig. 2a). Similarly, during fasting, alpha diversity (Shannon diversity) was significantly reduced relative to baseline, which also normalised during follow up (Fig. 2b). To contextualise fasting-induced gut microbial changes and compare their magnitude to other interventions or normal temporal fluctuation, we used longitudinal metagenomic data from five external studies, including another fasting intervention, an antibiotic intervention, a dietary intervention with whole grains and two observational studies without interventions (Fig. 2c). This showed that the immediate effects of fasting on the gut microbiome were substantially larger than observed in the whole grain intervention, but smaller than changes during antibiotic treatment (Fig. 2c). In stark contrast to the antibiotic cocktail, fasting did not have lasting long-term effects on the gut microbiome, as the differences to baseline at the follow-up timepoints (30-90 days) were similar to those observed for comparable time periods in the studies without any intervention. When resolving the effects of fasting at the genus and species level, we observed significant differences in the relative abundances of 66 out of 100 genera tested (42 increased, 24 depleted, FDR < 0.05 using linear mixed models (LMMs), see Fig. 2d and table S3). Interestingly, while in some genera all species displayed a similar direction of change (e.g. within *Lachnoclostridium*, *Actinomyces*, *Faecalibacterium* and *Roseburia*) in other genera changes were rather disparate and appeared to be species-specific (e.g. within *Bacteroides* and *Blautia*). At the level of individual bacterial species, fasting significantly affected the relative abundance of 516 out of the 772 bacterial species tested (306 depleted, 210 increased, FDR < 0.05 using LMMs, Fig. 2d and Table S4). For *Akkermansia muciniphila*, which is often postulated to mediate beneficial effects of fasting^31^, we observed large individual-specific changes, but across all participants this was not significant (adjusted p-value = 0.10, Extended Data Fig. 1). *Clostridioides difficile*, which was previously reported to outgrow during a very low calorie diet^32^, was not detected in a single sample from our large patient population.

**Figure 2:**
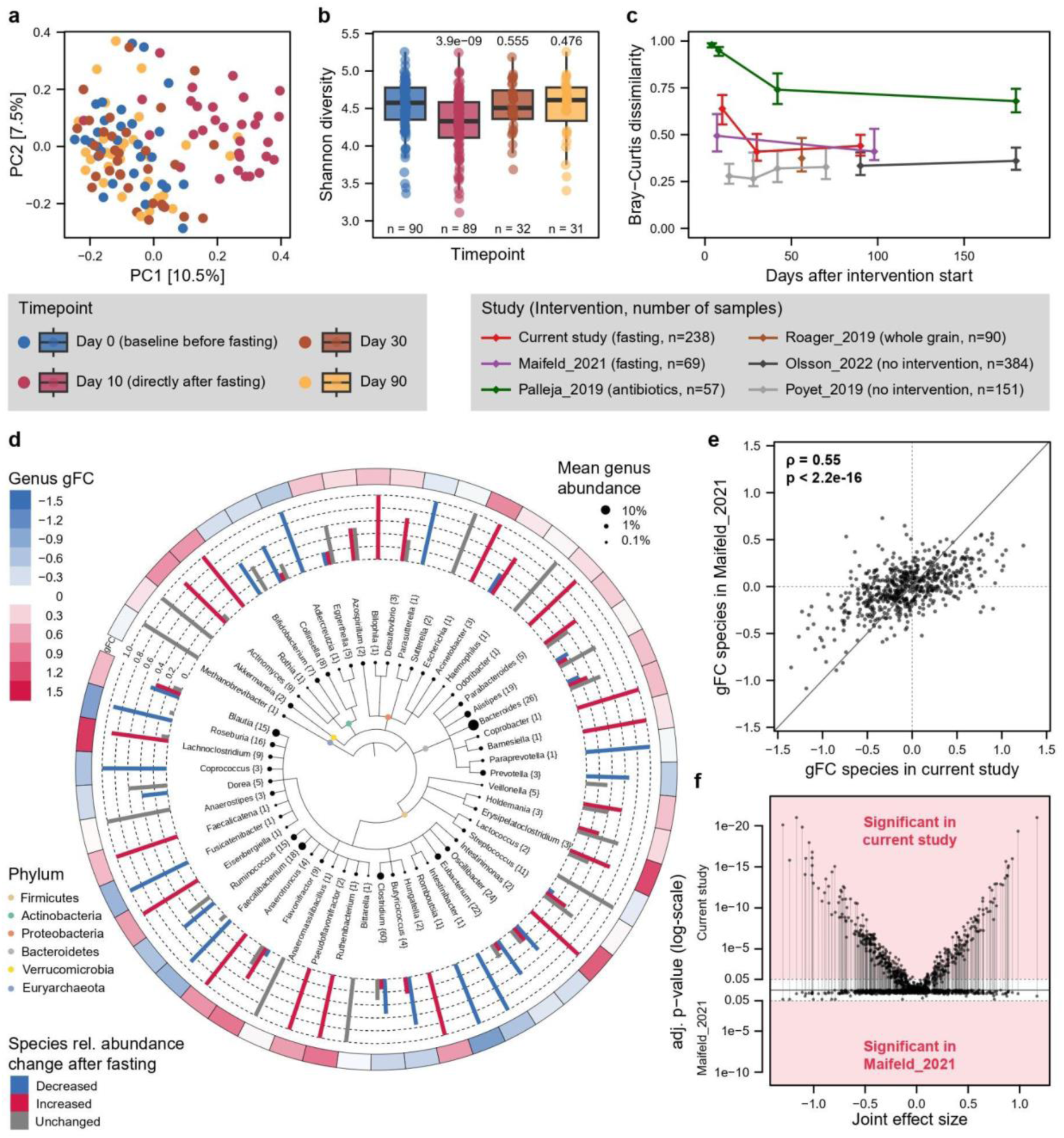
Changes at the taxonomic level in the gut microbiome during fasting. **(a):** PCoA on Bray-Curtis dissimilarities computed on all individuals (n = 28) from whom faecal samples were collected at all four timepoints. **(b)**: Shannon diversity of all samples (n = 241) over the four timepoints with significance tested using LMMs with a random effect per individual. Boxplots are defined as in Fig. 1. **(c)**: Longitudinal fluctuation of the gut microbiome as measured by Bray-Curtis dissimilarities within individuals over time relative to baseline for our study in comparison to other longitudinal studies with varying interventions. Median is indicated by diamonds, while vertical bars correspond to the 25th and 75th percentile. **(d)**: Taxonomic tree showing the magnitude of change for prevalent genera (present in >40% of samples) and the directionality of change during fasting for all species within each genus. Numbers in curly brackets indicate the number of species within a given genus that have been tested for differential abundance (Methods). **(e)**: Validation of observed taxonomic changes by correlating fasting effects observed here (and quantified by the generalised fold change (gFC)) for each species to the data by Maifeld et al. using Spearman correlations. Only species occurring in both datasets and meeting the prevalence threshold in both studies are included **(f)**: Volcano plot of differential abundance of species in Maifeld et al. and the current study. Generalised fold changes were calculated using LMMs with a random intercept for both the individual and the study, while adjusted p-values were calculated in a per-study fashion using LMMs with a random intercept for the individual.

We compared our results to the only other published study which has used shotgun metagenomics to investigate the effects of fasting^1^, and found that abundance changes were generally in agreement for species detected in both studies (Spearman’s rho = 0.55 and p < 2.2e-16, Fig. 2e). It should be noted though that while no species reached the significance threshold in Maifeld et al. with the established statistical methodology used here, many of their largest reported changes were concordant with, and significant in, our study (Fig. 2e,f). Lastly, consistent with the beta-diversity analysis mentioned above, we did not observe significant taxonomic differences at both the one and three-months follow-up compared to baseline, (no differentially abundant species at both long-term follow-up timepoints detected with adjusted p-values < 0.3). Altogether, our results show that fasting causes the majority of microbial species in the gut to substantially change in relative abundance, with a quick restoration to prior configuration after fasting cessation and resumption of food intake.

### Fasting remodels the functional capacity of the gut microbiome

Given the large taxonomic changes observed and considering that (almost) no nutrients reach the large intestine during fasting, we next aimed to understand the functional consequences at the level of microbial metabolism. To this end, we investigated changes in metagenomic gene abundances summarised at several functional levels with a focus on gut metabolic modules (GMMs) and CAZymes. Surveying high-level metabolic processes defined by the GMMs, we observed a significant increase in, amongst others, glycoprotein degradation and depletions in amine, polyamine and (complex) carbohydrate degradation during fasting (FDR < 0.05 using LMMs, Fig. 3a and Table S5). Similar to taxonomic changes, at the follow-up timepoints, no significant differences were observed relative to baseline. We further investigated whether a previous finding that urea nitrogen recycling by gut microbes contributed to protein balance in hibernating thirteen-lined ground squirrels, also applied to fasting humans^33^. To this end, we calculated differential abundances of all KEGG KOs in the urease operon as described previously^33^, but could not observe an increase in this operon. In contrast, 5 out of 7 KOs were depleted during fasting (Extended Data Fig. 2), which suggests that gut microbiome-mediated urea nitrogen recycling does not contribute to protein balance in fasting humans.

**Figure 3:**
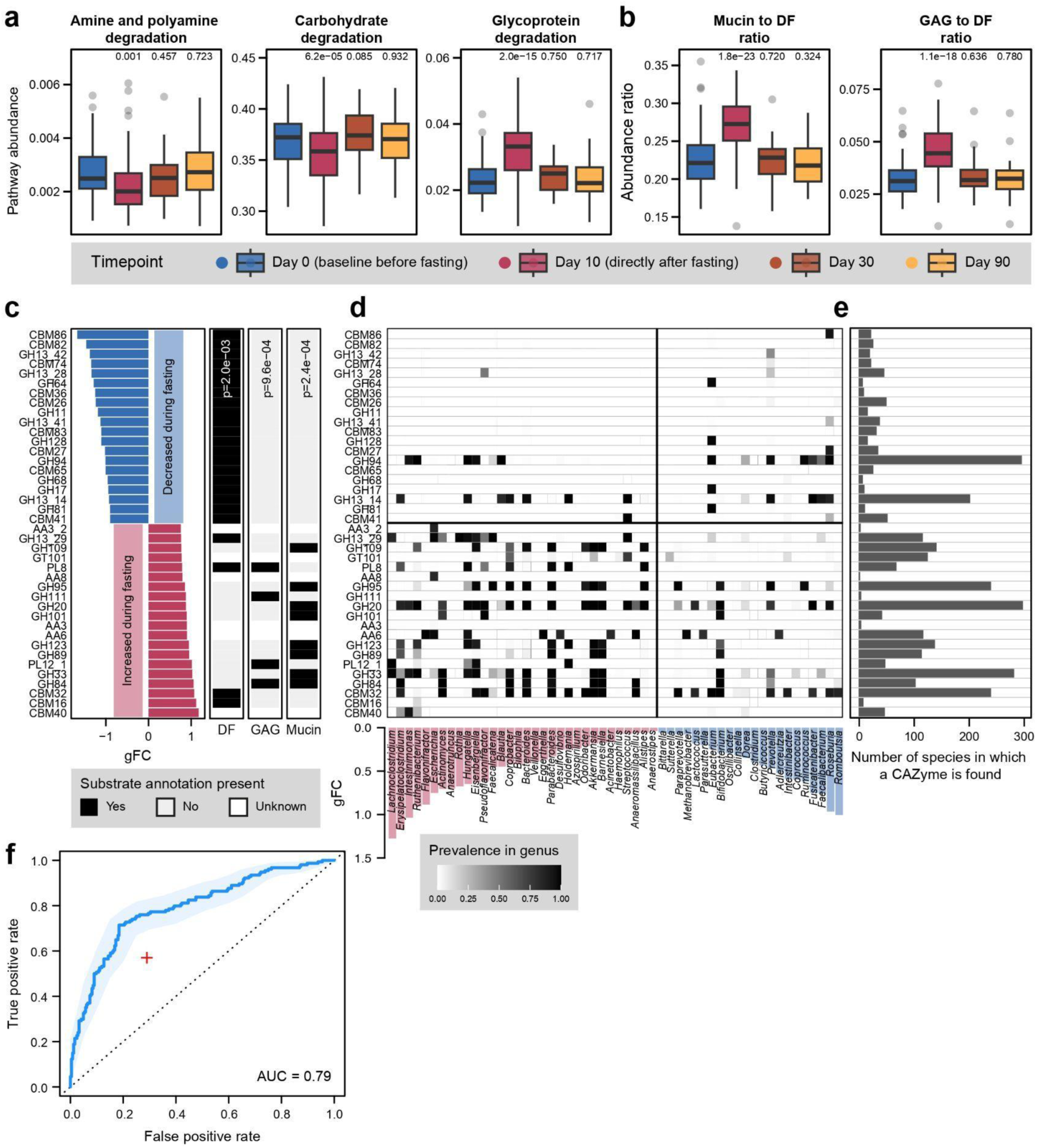
Metagenomic remodelling of gut microbial energy metabolism during fasting. **(a):** Changes in relative abundance of gut metabolic modules (GMMs) across the study period. **(b):** Boxplots of the ratio between the total abundance (RPKM) of mucin-targeting and dietary-fibre targeting CAZymes and between the total abundance of glycosaminoglycan(GAG)-targeting and dietary-fibre targeting CAZymes. **(c):** Bar plots showing the differential abundance of the 20 most enriched CAZymes during fasting (red) and the 20 most depleted CAZymes during fasting (blue) based on generalised fold changes (all 40 displayed CAZyme values have adjusted p-values < 1e-5). All CAZyme families were additionally subjected to gene-set enrichment analysis (GSEA) to pinpoint substrate enrichments (GSEA p-values are indicated underneath). **(d):** Heatmap indicating the genomic prevalence of the increased and decreased CAZymes in genera (Methods). Genera are coloured based on whether they significantly increased (left half, red vertical bars) or decreased (right half, blue vertical bars) during fasting (Fig 2D and generalised fold changes are indicated). **(e):** Number of species that genomically encode a given CAZyme (rows corresponding to (c)). All species included in differential abundance analysis were surveyed. **(f):** ROC curve for a classifier cross-validated to predict if a species increases or decreases in relative abundance during fasting based on the genomic CAZy repertoire of a given species (see inset for AUC) and blocked on the taxonomic family level. Red cross indicates performance of a classifier using genomic similarity between species (Methods). Boxplots are defined as in Fig. 1. Significance for all box plots is calculated by employing linear mixed models with the participant modelled as a random effect.

Next, to survey preferential carbohydrate substrate use, we computed high-level substrate ratios based on metagenomic CAZyme genes. This showed that during fasting there is a strong increase in mucin-to-dietary fibre (Fig. 3b) and glycosaminoglycan-to-dietary fibre ratios (Fig. 3b) consistently indicating increased microbial metabolism of host glycans. When surveying specific CAZyme families that are differentially abundant, we indeed saw that dietary fibre-metabolising CAZymes were strongly depleted during fasting, while CAZymes involved in mucin foraging and GAG metabolism were strongly increased (Fig. 3c and Table S6). In order to understand the taxonomic origin of differentially abundant CAZymes, we annotated the CAZyme repertoire in 107,683 bacterial gut genomes. We then computed for each genus the mean prevalence of each individual CAZyme (Methods) and plotted the genomic CAZyme repertoires of the most strongly depleted / increased genera, which showed concordance with CAZyme enrichments (Fig. 3c,d). In order to gain more insight into the taxonomic spread of a CAZyme family and how many species potentially contribute to the abundance of a given CAZyme, we also counted the number of species in which each CAZyme occurred (Fig. 3e).

Lastly, given the strong substrate switch in microbial metabolism, we wondered whether we could predict if a specific bacterial species would increase or decrease in relative abundance during fasting based on their genomic CAZyme repertoire. Indeed, this was possible with high accuracy (average area under the receiver-operating characteristic curve [AUC] of 0.79, see Fig. 3f), even though taxonomic relatedness itself was also predictive of relative abundance changes, albeit with lower accuracy (Fig. 3f) (Methods). Altogether, fasting-induced alterations in the gut microbiome were not only clearly delineated by functional metagenome analyses, but also readily interpretable in terms of shifts in energy substrate utilisation. Indeed, fasting-induced taxonomic changes could be predicted from the genomic repertoire of the respective bacterial taxa.

### Impact of fasting on the serum metabolome and inference of bacterial producers

We next investigated longitudinal serum metabolomic profiles (either 3 or 4 timepoints, Fig. 1b) in 66 participants, for whom a total of 230 serum metabolomes with 721 unique metabolites had been measured (Table S7 for normalised metabolite abundances of all samples). We observed 382 out of 721 metabolites to be significantly changed directly after fasting (Day 10 at an FDR < 0.05 using LMMs), of which 246 metabolites were increased and 136 decreased (Table S8). As expected, a strong increase during fasting was observed for the ketone bodies acetoacetic acid and 3-hydroxybutyric acid (Fig. 4a-c). Other strongly increased metabolites included L-homocysteic acid and 2-hydroxybutyric (Fig. 4b,c), which likely reflected protein usage during fasting for gluconeogenesis. This protein source was not necessarily muscle-derived, as muscle has protein-sparing mechanisms in place, but likely coming from other tissues undergoing autophagy or degradation of the extracellular matrix^34^. On the other hand, among the most depleted metabolites were theophylline and the secondary bile acids taurodeoxycholic acid and litocholic acid (Fig. 4a). Theophylline (1,3,7-trimethylxanthine) is a commonly used drug for treating asthma and chronic obstructive pulmonary disease, but we believe this depletion rather reflected a decreased nutritional intake, as it can be found in low amounts in teas, coffee and chocolate, amongst other dietary sources. Similarly, we observed large depletions in trigonelline and caffeine (1,3-dimethylxanthine).

**Figure 4:**
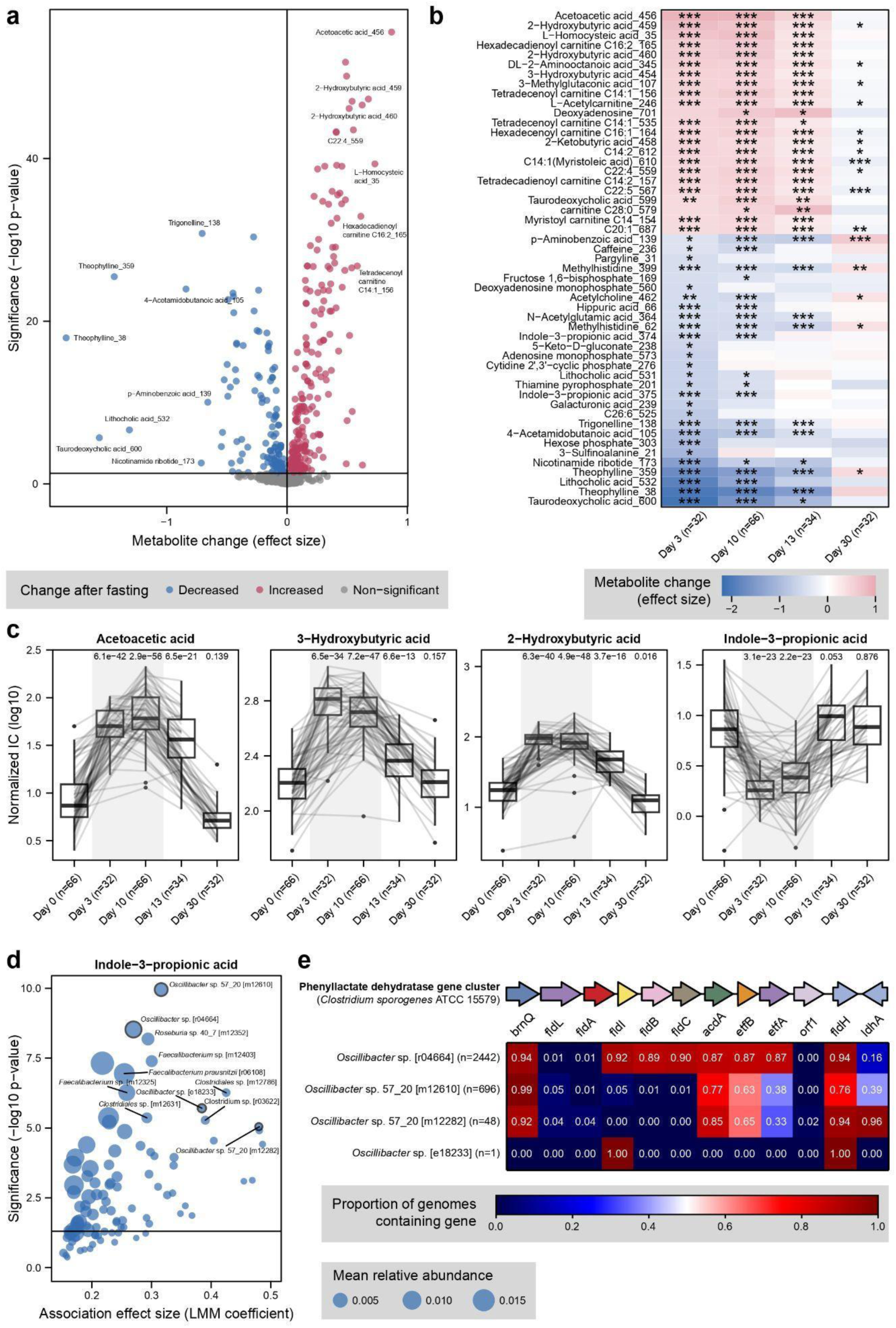
The switch in gut microbiome metabolism during long-term fasting impacts serum levels in metabolites of microbial origin. **(a):** Volcano plot showing the LMM estimates and statistical significance of the 721 metabolites changing in abundance during fasting (total n = 132 samples). Metabolites with an FDR-corrected p-value < 0.0001 and effect size > 0.25 or < -0.25 are labelled. **(b)**: Heat map showing the time-dependent changes in abundance during and directly after fasting as compared to baseline (n = 66). Metabolites are ordered based on their mean effect size estimate for Day 3 and Day 10, as samples on these timepoints were obtained during and at the end of fasting. Only metabolites with an effect size > 0.5 or < -0.5 and an FDR-corrected p-value < 0.05 at Day 3 or Day 10 are shown. FDR-corrected P-values are indicated by * < 0.05, ** < 0.001, *** < 0.0001 **(c)**: Time trajectory for the abundance of 4 significantly different metabolites (low p-value, high effect size) with each grey line representing one study participant. IC: Ion Current **(d)**: Volcano plot showing LMM estimates and statistical significance for the intra-individual association between indole-3-propionic acid (IPA) and bacterial species abundance. Significance and enrichment effect sizes were obtained with LMMs (modelling metabolite levels using species relative abundance as predictor, with participant as random effect) and only species > 30% prevalence were included in the models (Methods). Dot size corresponds to the mean relative abundance of the respective species at baseline (n=90 samples) **(e):** Proportion of genomes per *Oscillibacter spp.* that contain genes homologous to those in the phenyllactate dehydrogenase gene cluster responsible for IPA production in the model organism *Clostridium sporogenes* (Methods). Number of genomes screened per species is indicated in brackets.

Among the decreasing metabolites we also noted indole-3-propionic acid (IPA), a microbial product of tryptophan degradation. Curiously, few gut bacteria (*Clostridium sporogenes,* some other *Clostridium* spp., and several *Peptostreptococcus* spp.) have been described to have the metabolic capacity for this conversion^35^. Importantly, most of these species were not detected at all or at very low prevalence in our study or external large cohorts, indicating that they are unlikely to be important IPA producers in the human gut (Extended Data Fig. 3). To identify additional potential producer species of this metabolite, we employed LMMs to identify species linearly co-varying with IPA abundance over time (Fig. 4d). This yielded several species with significant intra-individual associations to IPA levels, with four *Oscillibacter* species among these (Fig. 4d). As expected based on their temporal association with IPA, all four *Oscillibacter* spp. decreased in abundance during fasting (Extended Data Fig. 4). Moreover, these four *Oscillibacter* spp. also exhibited strong inter-individual correlation with IPA abundance at baseline (Spearman’s rho between 0.27 and 0.45, all p-values < 0.05, Extended Data Fig. 4). When we investigated public *Oscillibacter* genomes for the presence of genes homologous to the 15-kb phenyllactate dehydratase gene cluster known to be responsible for the conversion of tryptophan to IPA, as described in *Clostridium sporogenes*^36^, we found strongly homologous genes in genomes of two out of the four Oscillibacter species (Fig. 4f) (Methods). Compared to the gene cluster described in *C. sporogenes*, the *Oscillibacter* spp. often lacked *fldL*, mimicking the gene arrangement in several *Peptostreptococcus* species that were found to produce IPA despite lacking *fldL*^37^. As *fldA* (an acyl-CoA transferase) is also often lacking in *Oscillibacter* spp., we speculate that this role can be taken over by distantly homologous genes capable of acyl-CoA transfer. In summary these results suggest *Oscillibacter* species to be prevalent and abundant producers of IPA in the human gut (Extended Data Fig. 3).

## Discussion

Longer periods of food shortage have always been a common occurrence during human evolution, until very recent centuries. While the human physiological responses to extended periods of fasting and caloric restriction have been well-studied, adaptations of the gut microbiome to this situation have remained largely elusive. In this work we provide a detailed picture of (functional) microbiome responses to fasting and link this to serum metabolites in the largest study of its kind to date. By further performing bacterial genome mining to discover producers of microbial metabolites, we uncover *Oscillibacter* species likely capable of IPA production. Due to their high prevalence and abundance, we hypothesise these to be the main bacterial producers of IPA in the gut. IPA is a microbial metabolite with strong clinical relevance for cardiometabolic diseases^38,39^ and molecular mechanisms have been described through which IPA improves cardiometabolic health. For example it protects against diastolic dysfunction by enhancing the nicotinamide adenine dinucleotide salvage pathway^40^ and inhibits atherosclerosis development by facilitating macrophage reverse cholesterol transport^39^. Unfortunately, very few *Oscillibacter* isolates are available for experimental validation of our finding, likely due to difficulties in culturing isolates from this genus^41^. Our results however suggest that enhancing fasting with prebiotic supplementation aimed at maintaining or increasing *Oscillibacter* abundance might further improve (cardiometabolic) health benefits.

To obtain a detailed view on substrate utilisation by gut microbes during fasting, we combined CAZyme and GMM analyses. While it was previously shown that mucin-foraging bacteria increase during long-term fasting^1,29^, this had not been resolved at the level of individual CAZymes. Curiously, one of the most well-known mucin foragers in the human gut, *Akkermansia muciniphila*, was not significantly affected by long-term fasting, neither in our study nor in re-analysed data of another one (Extended Data Fig. 1)^1^, which contrasts with many earlier reports of its enrichment during various forms of fasting^31,42^. Instead, in our study, we find other mucin-foraging bacteria to strongly increase in relative abundance during fasting, including *Ruminococcus torques* and *Ruminococcus gnavus*. Interestingly, this finding resembles ones from studies of fibre-deprived diets in mice, even though these animals still consumed macronutrients^43,44^. While increased mucin foraging has often been viewed as negative for human health due to possible mucus layer erosion^43,44^, the health implications of this in the context of fasting are less clear given that intestinal regeneration and improvement of the gut barrier take place during fasting^5^.

A main strength of our study is the joint collection of patient blood markers, metagenomic and metabolomic data in a longitudinal design with an unprecedentedly large participant population, surveyed under controlled clinical conditions. Major limitations of our study first relate to its design, precluding us to disentangle which beneficial metabolic health effects are mediated through the gut microbiome or occur independently of it. This could potentially be addressed in future studies that include an additional study arm which more directly perturbs the gut microbiome, e.g. through administration of antibiotics or dietary fibre. Despite this remaining uncertainty about causal effects, the remarkable congruency between our data and that of a previous study^1^ (Fig 2) rules out that the effects observed here are attributable to centre-specific or study-specific uncontrolled variables. A second limitation relates to the lack of granularity in our repeated sampling design, including only one sample at baseline, which prohibits the study of high-resolution temporal dynamics before as compared to during or right after fasting. In addition, the observed complete reversion to baseline microbiome composition as early as a few weeks after food reintroduction indicates that it would be more useful to include additional early time points rather than the three-month follow-up time point. As metabolic improvements due to fasting have been described to be long-lasting (although they partially depend on lifestyle after fasting)^45^, our observation that microbiome composition quickly returns to baseline, poses the intriguing question of whether and how (drastic) short-term microbiome changes could contribute to long-term improvement of metabolic health.

In conclusion, our study provides a view of unprecedented resolution about functional gut microbiome responses to long-term fasting and how these are associated with changes in serum metabolite levels. This raises interesting possibilities for the development of future fasting or dietary intervention studies with the dedicated aim of exploring and maximising microbiome-dependent health benefits.

## Data Availability

Raw sequence data of the current study will be released upon acceptance of the manuscript under accession number PRJEB74035 at the European Nucleotide Archive (ENA). Sequence data from external cohorts used in this manuscript can be found under the following accession numbers: PRJNA698459, ERP022986, PRJNA395744, PRJEB38984, SAMN11950000–SAMN11950562, PRJEB39223 and PRJEB11532. All processed profiles and files can be accessed through https://github.com/zellerlab/Buchinger_Long_Term_Fasting.

## Code Availability

All custom code and required data files to reproduce the results and figures can be found at https://github.com/zellerlab/Buchinger_Long_Term_Fasting.

## Acknowledgements

We thank members of the Zeller group for fruitful discussions. We are moreover indebted to the EMBL IT Services Team for support with high-performance computing. This work received funding from EMBL, the Buchinger Wilhelmi Development & Holding GmbH, the Federal Ministry of Education and Research (BMBF grant no. 031L0181A to G.Z.), the German Research Foundation (Deutsche Forschungsgemeinschaft project number 395357507 – SFB 1371 to G.Z.), the LUMC (LUMC Fellowship to G.Z.), the Health + Life Science Alliance Heidelberg Mannheim through state funds approved by the State Parliament of Baden-Württemberg (Postdoctoral Fellowships to Q.D. and N.K.) and an EMBO postdoctoral fellowship (EMBO ALTF 1030-2022 to Q.D.)

## Authors’ contributions

QD performed bioinformatic and statistical analysis and designed figures. FG, FWdT, CG and RM designed and performed the clinical study. FG and AL conducted the clinical study. MM and SM performed serum metabolomics measurements. NK, ML and SR aided in bioinformatic analyses. QD, GZ and RM wrote the manuscript. GZ and RM supervised this work.

