## Extended Data Figures for "Long-term fasting remodels gut microbial metabolism and host metabolism"

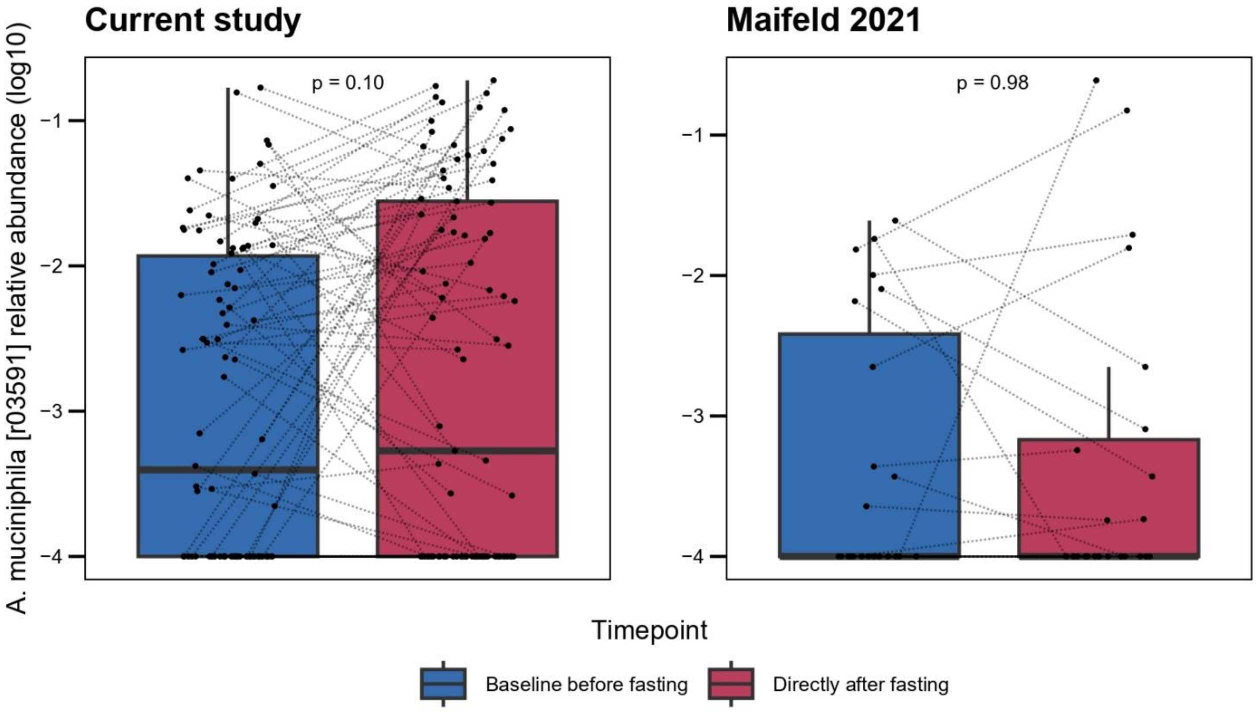


**Extended Data Fig 1: Relative abundance of *Akkermansia muciniphila* before and during fasting**. Dotted lines connect measurements of the same individual before and directly after fasting. The boxplot centre value corresponds to the median, the box indicates the interquartile range, and whiskers extend to 1.5 times the interquartile range. Significance is calculated by employing linear mixed models with the participant modelled as a random effect and p-values were adjusted using the FDR. Duration of fasting was slightly different between the current study (ten days) and Maifeld et al. (seven days).

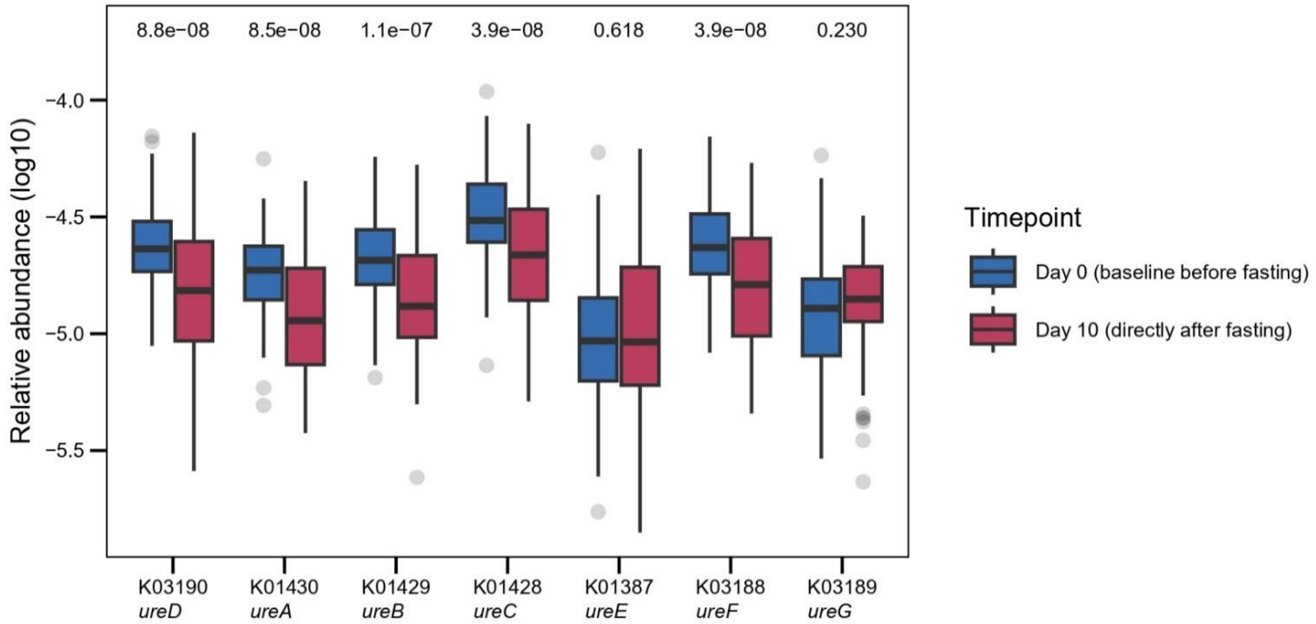


**Extended Data Fig 2: Abundances of urease operon before and during fasting.** Abundances of individual KEGG KOs contained in the urease operon[33]. Boxplots are defined as in Ext. Data Fig 1 with dots representing outliers. Significance is calculated by employing linear mixed models with the participant modelled as a random effect and p-values were adjusted using the FDR.

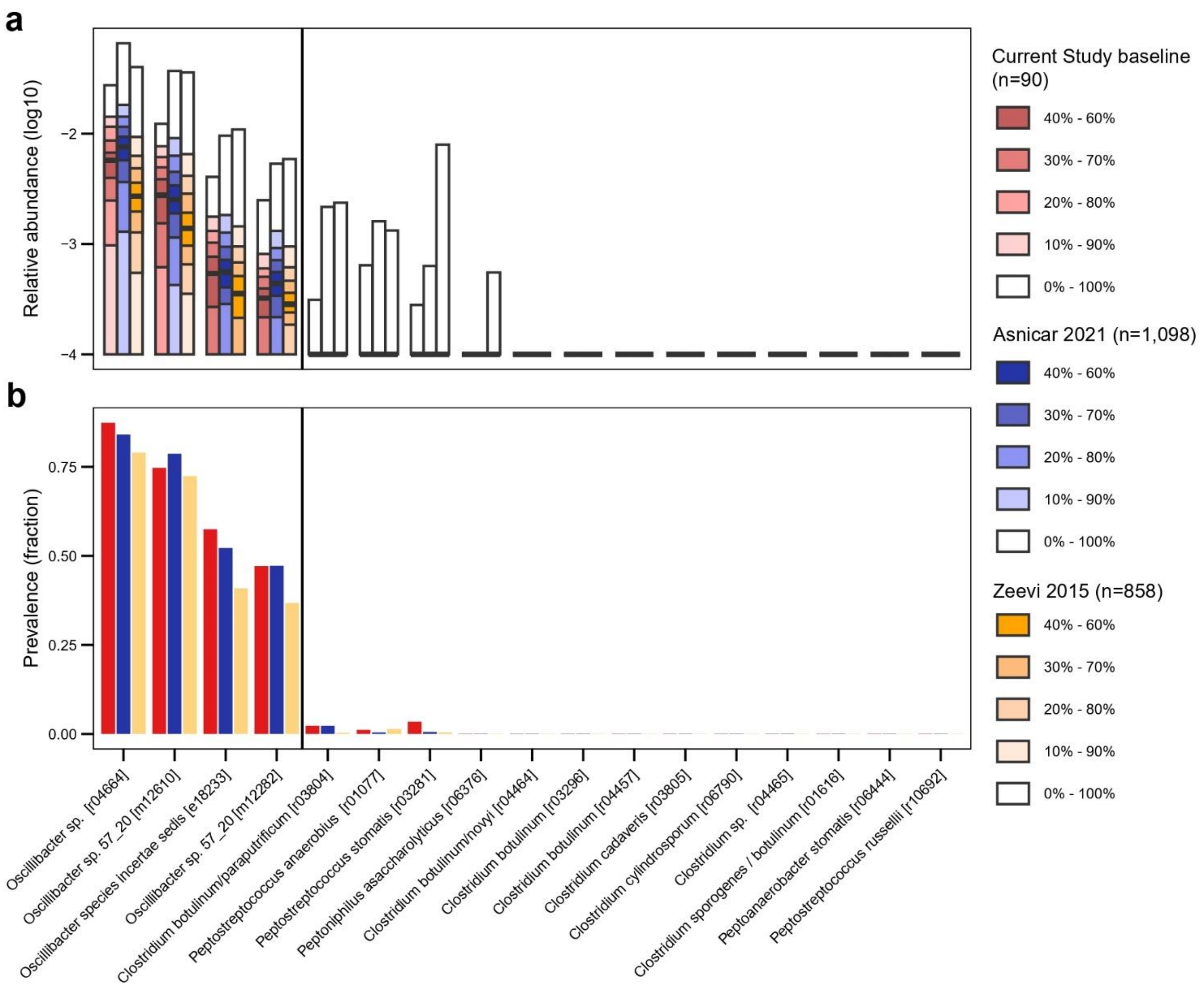


**Extended Data Fig 3: Relative abundance and prevalence of inferred IPA-producing gut bacteria across geographically diverse cohorts.** **(a)** Relative abundance of the four *Oscillibacter spp.* associated with IPA levels as presented in Fig. 4d alongside known IPA-producing gut bacterial species from literature, displayed as quantile plots in baseline data from the current study (n=90) and large-scale metagenomic cohorts from the Zeevi et al. (n=858) and Asnicar et al. (n=1,098) studies[19,20]. A pseudo-count of 1e-4 was added for visualisation purposes. **(b)** Prevalence of the same species across the respective cohorts. Rarefaction was performed prior to prevalence calculation (Methods). This resulted in slightly lower sample numbers (n=87, n=851 and n=1093 for the baseline of the current study, Zeevi and Asnicar, respectively).

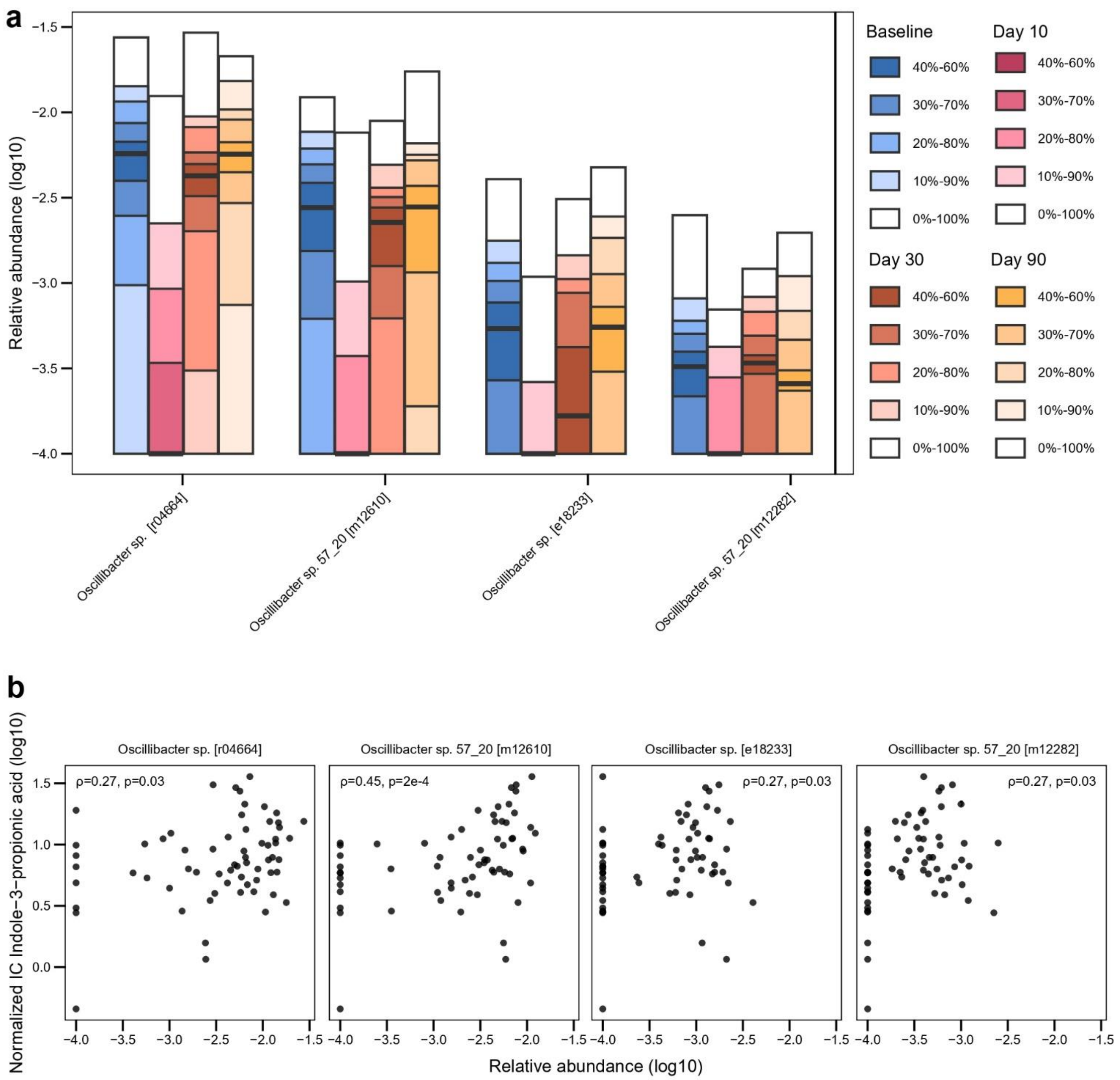


**Extended Data Fig 4**: **Relative abundances of *Oscillibacter* spp. and correlations with IPA levels**. **(a)** Relative abundance of the four *Oscillibacter* spp. associated with IPA levels as shown in Fig. 4d across all time points (n=90 at baseline, n=89 during fasting (Day 10), n=32 at Day 30 and n=30 at Day 90) of our study. Data is displayed as quantile plots[21]. **(b)** Spearman correlation between IPA abundance and the four *Oscillibacter* spp. across the participants in our study at baseline (n=64 samples with matching serum metabolomics and faecal metagenomics) as identified in the LMM analysis shown in Fig. 4d.
